# The infectious unit of influenza A virus is a population of particles, not a single virion, as shown by absolute particle quantification

**DOI:** 10.64898/2026.11.11.765807

**Authors:** Yuu Kawahara, Kenji Tamao, Yoichiro Fujioka, Seiya Yamayoshi, Yusuke Ohba, Yoshihiro Kawaoka, Hiroyuki Noji, Kazuhito V. Tabata

## Abstract

Influenza A virus (IAV) populations are characterized by a high prevalence of non-infectious virions, with particle-to-plaque forming unit (PFU) ratios typically ranging from 10 to 1,000. While “multiplicity reactivation“—the process by which incomplete viral genomes (IVGs) establish productive infection through co-infection—has long been hypothesized, its quantitative nature has remained elusive. In this study, we utilized a digital influenza assay to achieve absolute quantification of IAV particles, allowing us to define the particle-based multiplicity of infection (pMOI) for precise evaluation. We combined this approach with a single-cell infection assay using microcompartmentalized containers and machine learning-based automated cell classification to accurately assess infection outcomes across a wide range of pMOIs. Our analysis of the A/Puerto Rico/8/1934 (H1N1) revealed a count-to-plaque forming unit (PFU) ratio of approximately 400, meaning only a tiny fraction of particles possesses infectivity. Remarkably, the infection rate in single-cell infection experiments followed a sigmoidal curve and reached saturation at a pMOI of only 5 to 10. This demonstrates that under high-pMOI conditions, productive infection is predominantly driven by genome complementation among incomplete particles rather than solely by fully infectious virions. On the other hand, mathematical model analysis based on these findings suggested that the functionality of each of the eight genomic segments within a virus particle follows a stochastic process, resulting in a probability distribution for the number of functional segments per particle. This mechanism proved exceptionally robust across multiple strains despite a 20-fold variation in CTPR values. Ultimately, these findings indicate that incomplete particles are not merely biological noise but active contributors to the infection process, significantly enhancing the infectivity and evolutionary flexibility of IAV. This study will provide a more refined understanding of influenza transmission dynamics and risk assessment, particularly in environments where multiple virus particles are present within a single respiratory droplet.

## Introduction

Influenza causes annual epidemics, affecting an estimated one billion people worldwide each year. Furthermore, as it repeatedly causes pandemics, it has garnered global attention as a significant public health issue. One reason for these recurring outbreaks is the high mutation rate of the influenza virus. The mutation rate of influenza A virus (IAV) is as high as 10^−4^ substitutions per nucleotide per replication cycle^1^. Additionally, the segmentation of the IAV genome into eight segments is a key driver of its rapid evolution. When two distinct IAV strains co-infect a single cell, progeny viruses are produced as chimeras containing mixed combinations of genome segments from both parent particles. This phenomenon, known as reassortment, is believed to be closely linked to host adaptation and the emergence of novel viruses ^2–4^. Consequently, the co-infection of individual cells by multiple virions is likely to play a critical role in viral evolution.

Following entry into a host cell, virus particles release eight genome segments, which are replicated to produce progeny virions. This implies that viral reproduction may fail if all eight genome segments are not successfully delivered into the cell ^5^. On the other hand, the particle-to-PFU ratio of IAV is typically between 10 and 1,000 ^6–8^. This indicates that the vast majority of virions in a population are non-infectious. Furthermore, single-cell transcriptomic analyses have revealed highly heterogeneous expression of viral mRNAs and proteins, including the complete absence of certain segments in many infected cells ^9–12^. Previous studies using electron microscopy have shown that the proportion of purified virions containing all eight segments ranges from 20% to 80% ^13,14^. These observations suggest that cells infected at a low multiplicity of infection (MOI) consist of those with lack of one or more genome segments or non-functional segments, meaning the majority of IAV particles function as incomplete viral genomes (IVGs). Consequently, only a fraction of the delivered segments ultimately functions within the cell. This implies that a viral population incapable of producing progeny from a single virion would face extinction as a collective, thereby highlighting the absolute necessity of a collective infection strategy for viral maintenance.

In IAV populations where incomplete particles predominate, co-infection of a single cell is critically important for achieving productive infection. This phenomenon, which forms the core of the reassortment concept proposed in the 1950s and 1960s, is termed “multiplicity reactivation” ^15–17^. This concept is based on the observation that when cells are inoculated with viruses whose genomes have been damaged by gamma rays or UV light, the hemagglutinin titer in the culture supernatant rises more rapidly in a manner dependent on the amount of virus inoculated, effectively restoring infectivity. Since then, discussions have been framed within the context of complementary infection by incomplete or partially damaged particles ^18–23^. However, these investigations have relied primarily on infectious titers or bulk RNA quantification (e.g., qRT-PCR), and research involving direct measurement of absolute virion numbers has been lacking. This represents a significant limitation, as particle count is the fundamental factor in multiple infection scenarios. Consequently, the true quantitative nature of co-infection has remained unclear.

Meanwhile, many studies quantify influenza viral load using plaque assays, which are the most widely used method, or molecular approaches such as qRT-PCR; however, these provide indirect measurements or capture only a subset of virions rather than providing direct particle counts. Furthermore, measurements of infectious titers only capture virions that successfully establish infection, which does not represent the total number of particles in a population containing non-infectious virions. The difficulty in quantifying total virus particles arises because individual virions lack independent growth activity and are too small to be observed with an optical microscope. Direct quantification requires specialized techniques, such as counting via electron microscopy ^24^, flow cytometry-based assays ^25^, or chromatographic analysis ^26^. As such, these methods are not easily accessible and have not been widely integrated into routine influenza virology research.

Recently, a “digital influenza assay” was developed to facilitate the simple and absolute quantification of IAV particles ^27^. Similar to digital PCR, this method stochastically captures and detects individual virions within micro-spaces. The digital influenza assay utilizes an array of approximately 50-fL microchambers to stochastically encapsulate particles and detects them via the fluorescence generated by their neuraminidase activity. In other words, the number of fluorescent chambers corresponds to the number of virus particles; thus, the concentration can be determined by normalizing this count by the total assay volume. This enables the easy quantification of virus particle concentrations in samples.

In this study, we define virus particle concentrations using the digital influenza assay and perform infection experiments based on the particle-based multiplicity of infection (pMOI) to quantitatively evaluate multiplicity reactivation. We determined the ratio of incomplete to complete particles in an Influenza A/Puerto Rico/8/1934 (H1N1) (PR8) preparation by comparing absolute particle counts from the digital assay with infectious titers from plaque assays. The results revealed an approximate ratio of 400 incomplete particles for every single infectious particle. We subsequently conducted infection rate measurements; however, standard plaque assays are limited at high MOIs because plaques overlap, making accurate counting impossible. To overcome this, we developed a single-cell infection assay to distinguish infection at the individual cell level. Using this system, we performed infection experiments across various pMOIs. The results showed that the infection rate reached its maximum at a pMOI of approximately 5. This implies that under high-pMOI conditions, the majority of infections are established by incomplete particles rather than fully infectious virions. Furthermore, experiments with viral samples exhibiting a difference in the ratio of incomplete to complete particles of over three orders of magnitude showed that infection saturation occurred at a pMOI of at most around 10. These findings demonstrate that multiplicity reactivation occurs even when the proportion of incomplete particles is extremely high, suggesting that infection via complementation is a general phenomenon in high-pMOI environments. Collectively, these findings suggest that the prevalence of incomplete particles in IAV does not impede population maintenance; rather, they actively participate in the infection process, thereby contributing to the high infectivity and evolutionary flexibility of the influenza A virus.

## Results

### Quantification of viral particles and infection titer

The influenza A/Puerto Rico/8/34 (H1N1) (hereafter referred to as PR8) was propagated in Madin-Darby canine kidney (MDCK) cells, and the number of virus particles and the infectious titer were measured. The particle number was determined using a digital counting method based on a microdroplet chamber^27^. In this method, influenza virus particles are probabilistically encapsulated in microdroplet chambers, and neuraminidase activity on the virus surface is detected using a fluorescent substrate. Therefore, the number of influenza virus particles can be quantified by counting the fluorescent chambers (Fig. 1A). The infectious titer was measured using a plaque assay with MDCK cells (Fig. 1B). The concentration of virus particles was measured to be 1.4(±0.4)×10¹¹ VP/mL, while the infectious titer was determined to be 3.6(±0.6)×10⁸ PFU/mL. Subsequent to the determination of these values, the count-to-PFU ratio (CTPR) was calculated to be 4.0(±1.3) ×10² VP/PFU (see Table 1). This indicates that a mere one in approximately 400 virus particles possesses the capacity for infection, thereby suggesting that the majority of the virus particles lack infectivity.

**Fig. 1.**
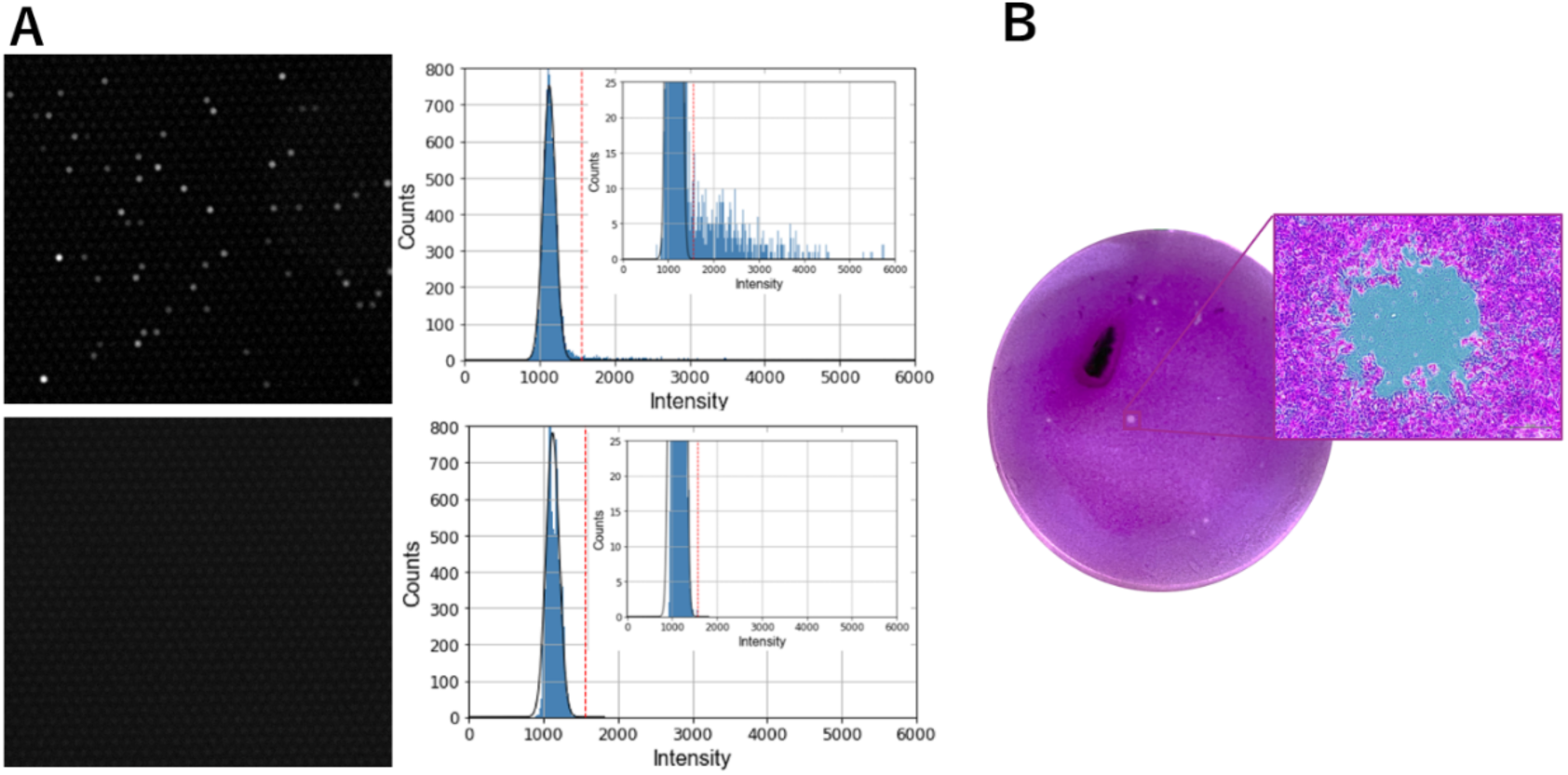
Quantification of influenza viruses by digital influenza assay and plaque assay. (A) Results of the digital influenza assay for PR8. The left panel shows the fluorescence images, while the right panel shows the distribution of fluorescence intensities, where the horizontal axis represents the fluorescence intensity of each chamber and the vertical axis represents the chamber count. The upper panels show the results from virus solution and the lower panels show the results of buffer solution. The red dashed line indicates the threshold defined as five times the standard deviation (5σ) of the background signal, and chambers with intensities above this value were counted as positive. The total count of positive chambers corresponds to the total number of virus particles, which was normalized by dividing by the total volume of the chambers to determine the virus particle concentration. (B) Results of the plaque assay for PR8.

**Table 1.** Ratio of Particle Number to Infectious Titer.

| Virus particles [VP/mL] | Virus titer [PFU/mL] | Count-to-PFU ratio(CTPR) [VP/PFU] |
|---|---|---|
| 1.4(±0.4)×10<sup>11</sup> | 3.6(±0.6)×10<sup>8</sup> | 4.0(±1.3)×10<sup>2</sup> |

**Table 2.** Ratio of Particle Number to Infectious Titer for Three Influenza Virus Strains.

| Strain | Virus particles [VP/mL] | Virus titer [PFU/mL] | Count-to-PFU ratio(CTPR) [VP/PFU] |
|---|---|---|---|
| H1N1 PR/8/1934 | 1.4(±0.4)×10<sup>11</sup> | 3.6(±0.6)×10<sup>8</sup> | 4.0(±1.3)×10<sup>2</sup> |
| H3N2 Aichi/2/1968 | 1.3(±0.4)×10<sup>11</sup> | 3.8(±0.5)×10<sup>7</sup> | 3.5(±1.1)×10<sup>3</sup> |
| H3N2 Texas/50/2012 | 8.5(±0.6)×10<sup>10</sup> | 9.2(±0.5)×10<sup>6</sup> | 9.2(±0.8)×10<sup>3</sup> |

### Measurement of infection rates under single-cell conditions

To achieve infection rate measurements under high-pMOI conditions, infection experiments were performed under single-cell conditions (Fig. 2A). Generally, the infection rate is calculated by dividing the number of infected cells (infection titer) by the total number of cells. On the other hand, the infection titer is usually determined by infecting cells cultured in a dish with a virus and observing the resulting cytopathic effect (CPE) or plaque formation. Therefore, under conditions of high MOI of 1 or more, all cells may be infected, which can lead to the formation of overlapping plaques or CPE. Thus, at high MOIs of 1 or more, it is difficult to accurately evaluate the relationship between viral load and infection rate. To address this, we added virus particles to a cell suspension containing a known number of cells at a pMOI calculated based on the number of virus particles, thereby inducing infection. We then encapsulated individual cells at sufficiently low concentrations in compartmentalized containers and observed infection at the single-cell level by monitoring cytopathic effects, such as cell burst and denaturation caused by viral release (Fig. 2A, B). In this experiment, the number of live cells was determined from cell images taken two days post-infection and demonstrated a decline in the number of live cells (Fig. 2C). The employment of machine learning algorithms enabled the automated classification of live cells within the captured images. The infection rates were calculated from the difference between the initial live cell count and the count of live cells two days post-infection. MDCK cells were infected with PR8 at various pMOI values, which were determined based on the number of viral particles measured by the digital influenza assay. On average, approximately 4.0×10³ cells were observed per condition, and their infection rates were measured (Fig. 2D, red dots). As a result, the infection rates followed a sigmoidal curve, beginning to rise from a pMOI of approximately 2. Furthermore, the infection rate reached approximately 80% at a pMOI between 5 and 10. Since the CTPR of the virus sample used in this study was 4.0 × 10², only one infectious particle capable of forming a PFU existed for every 400 particles. In other words, at a pMOI of 10, the probability of an infectious particle being present is only 2.5%. Moreover, if infection were caused solely by infectious particles, the infection rate would be expected to exhibit a linear increase, saturating only around a pMOI of 400. This result strongly suggested that under multiple infection conditions, infection is established even by incomplete particles such as defective viral genomes (DVGs) alone (Fig. 2E). These results allowed for the determination of the number of virus particles required to establish infection, revealing that the majority of these particles are DVG-like particles.

**Fig. 2.**
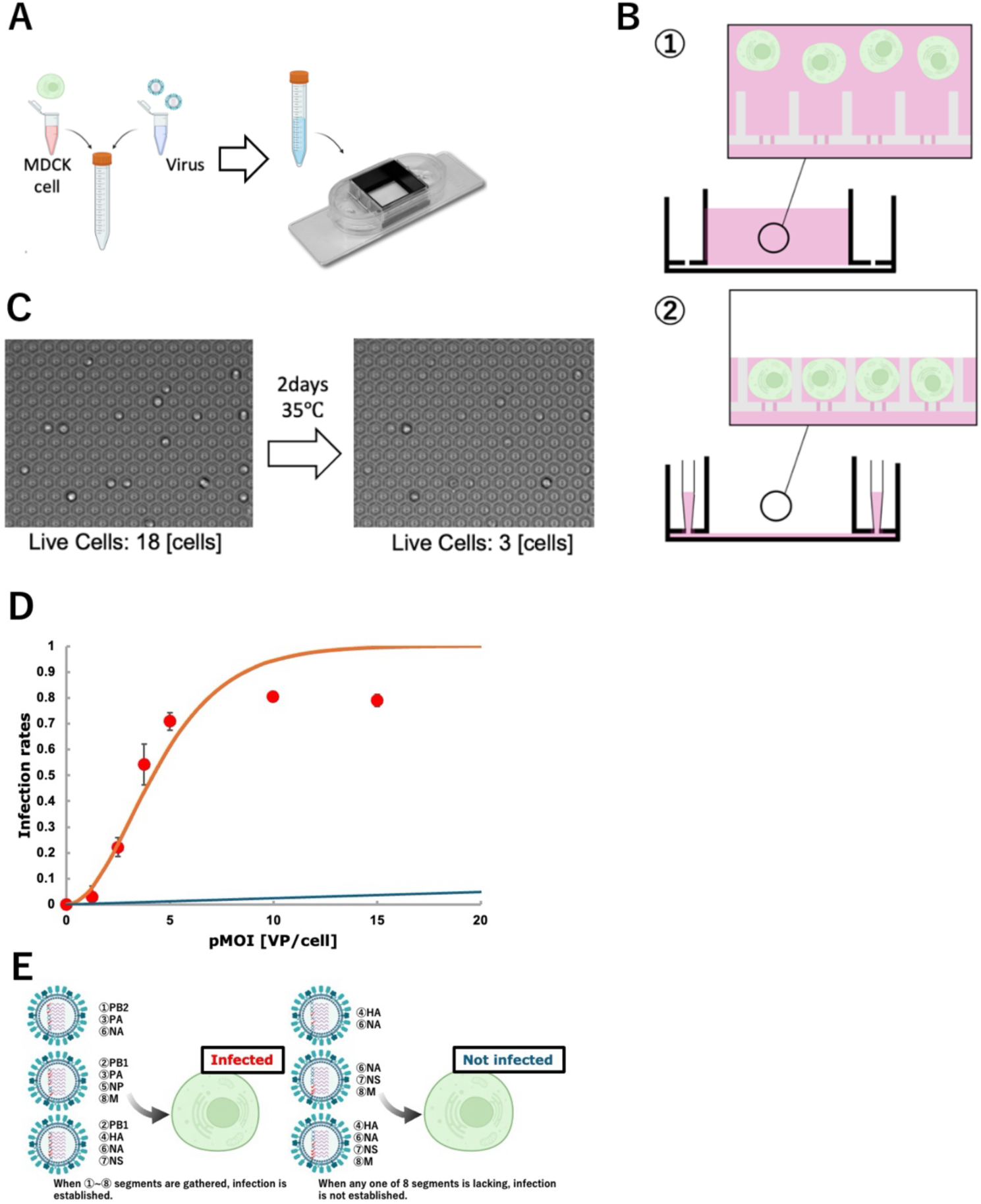
A measurement of infection rates under single-cell conditions and mathematical model of cellular infection. (A) Schematic diagram of the measurement method. (B) Diagram of the device used. The bottom of the microwells contains 2 µm holes, and cells are efficiently captured when solution is aspirated through side ports. (C) Cell images obtained immediately following infection and two days after infection. Several cells show degeneration due to viral infection. (D) Results of infection rate measurement under single-cell conditions and comparison with mathematical models. The x-axis indicates the mean number of virus particles per cell (pMOI), while the y-axis denotes the cellular infection rate. The red dots in the plot represent experimental data, the orange line represents the model with complementation, and the blue line represents the model without complementation. (E) Schematic illustration of the complementation and non-complementation model.

### Mathematical model assuming genome complementation

In order to better understand the experimental findings, a mathematical model of cellular infection was constructed. As demonstrated in prior research, the presence of DVG particles with partial deletions leading to the loss of genomic function has been substantiated^28,29^. Furthermore, electron microscopy observations of influenza virus particles have revealed that even wild-type virus populations contain particles missing one or more genome segments^5,13^. We developed a model assuming that DVG particles can complement each other’s missing genome segments during multiple infections and thereby establish infection. Initially, the probability that a virus particle contains an intact copy of each of the eight genome segments was calculated. Secondly, cellular infection rates were calculated based on the definition that infection is the gathering of eight intact genomic segments in a cell when several viruses invade the cell (Fig. 2E).

The probability that an intact genome segment is present in a viral particle was defined as the “Probability Present” (*P_P_*). We assumed that the virus particle count corresponds to the total number of virus particles, and that the infectious titer corresponds to the number of particles containing all eight intact genome segments. Therefore, the count-to-PFU ratio (CTPR) is indicative of the proportion of fully functional virus particles within the total viral population. *P_P_* (The probability of presence of a single intact genome segment) was calculated using the following formula (where *s* is the number of genome segments (for influenza virus, s = 8)):

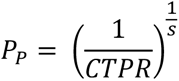

While the *P_P_* value of each genome segment may vary, previous RT-PCR studies have reported minimal differences among them^23^. Consequently, we hypothesized that there would be no correlation among genome segments and employed the geometric mean of the eight *P_P_* values as the representative *P_P_* for this study. Utilizing the CTPR value of 4.0×10² for the influenza virus sample (PR8), the calculated *P_P_* was 0.47.

Subsequently, we calculated infection rates under varying pMOIs. It is hypothesized that the *Vₙ* virus particles infect *Cₙ* cells (where *V*ₙ / *C*ₙ = pMOI), and that the number of viruses entering each cell follows a Poisson distribution. The probability of a cell being infected with genome complementation (*P_I_ (w/ complementation)*) was calculated by:

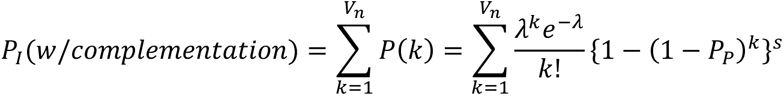

Conversely, in the absence of genome complementation, the probability of infection (*P_I_ (w/o complementation)*) was calculated simply based solely on the presence of fully functional virus particles.

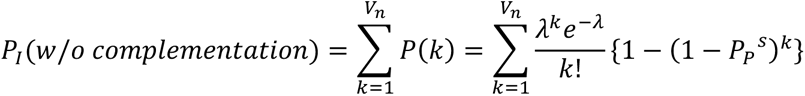

A comparison was made between the infection rates calculated by these two models and the experimental results from single-cell infection measurements (Fig. 2D; orange line: with complementation, blue line: without complementation). The model incorporating complementation (orange line) showed significantly higher infection rates from low pMOI compared to the model without complementation (blue line). Within the range of pMOI values that were examined, the infection rates in the no-complementation model remained at a very low level. This is due to the fact that the infection would rely solely on the extremely few fully functional virus particles. Additionally, the model assuming complementation exhibited a strong correlation with the experimental data (red dots). These findings suggest that incomplete genomes are complemented by functional genomes from other DVG particles during multiple infection, thereby enabling DVG particles to participate in productive infection. The favorable alignment between the theoretical model and the experimental results further suggests that the distribution of functional genome segments within virus particles adheres to a Poisson distribution.

### Measurement of infection rates at low pMOI using plaque assay

Under high pMOI conditions, a sigmoidal increase in the infection rate was observed, strongly suggesting that infection is established through the multiple infection of DVG particles. Conversely, under low pMOI conditions where multiple infection does not occur, a linear increase in the infection rate is expected, as infection would be caused solely by infectious particles that constitute PFUs. Therefore, to further validate multiple infection by DVG particles, we measured infection rates under low pMOI conditions using a plaque assay (Fig. 3A, red dots). In this experiment, the pMOI was set to a value significantly smaller than 1. Under such conditions, most cells remain uninfected, and it is a situation where a single virus particle rarely binds to a cell. Since multiple infection does not occur, complementation by DVG particles does not arise, and it is considered that only infectious particles can form plaques. Within the examined pMOI range, the model incorporating complementation (Fig. 3: orange line) showed a linearly increasing slope in the low pMOI region and an exponentially increasing slope as pMOI increased, whereas the model without complementation (Fig. 3: blue line) showed a straight line. While the difference in infection rates between the two models was small, the model incorporating complementation showed slightly higher values due to the rare stochastic occurrence of two-particle infections. Since infection at such low pMOIs depends on infectious particles containing all functional genome segments, a linear increase in infection rate was predicted. Experimental results showed that as pMOI increased, the observed infection rates deviated upward from the linear model without complementation and approached the model supporting complementation. This suggests the possibility of a complementary effect from two DVG particles that occurs stochastically and rarely even under low pMOI conditions, consistent with the model predictions. Nevertheless, under low pMOI conditions, the infection rate increased almost linearly, confirming that infection was driven primarily by infectious particles alone. This further demonstrates that under high pMOI conditions, complementation by DVG particles is the predominant cause of infection.

**Fig. 3.**
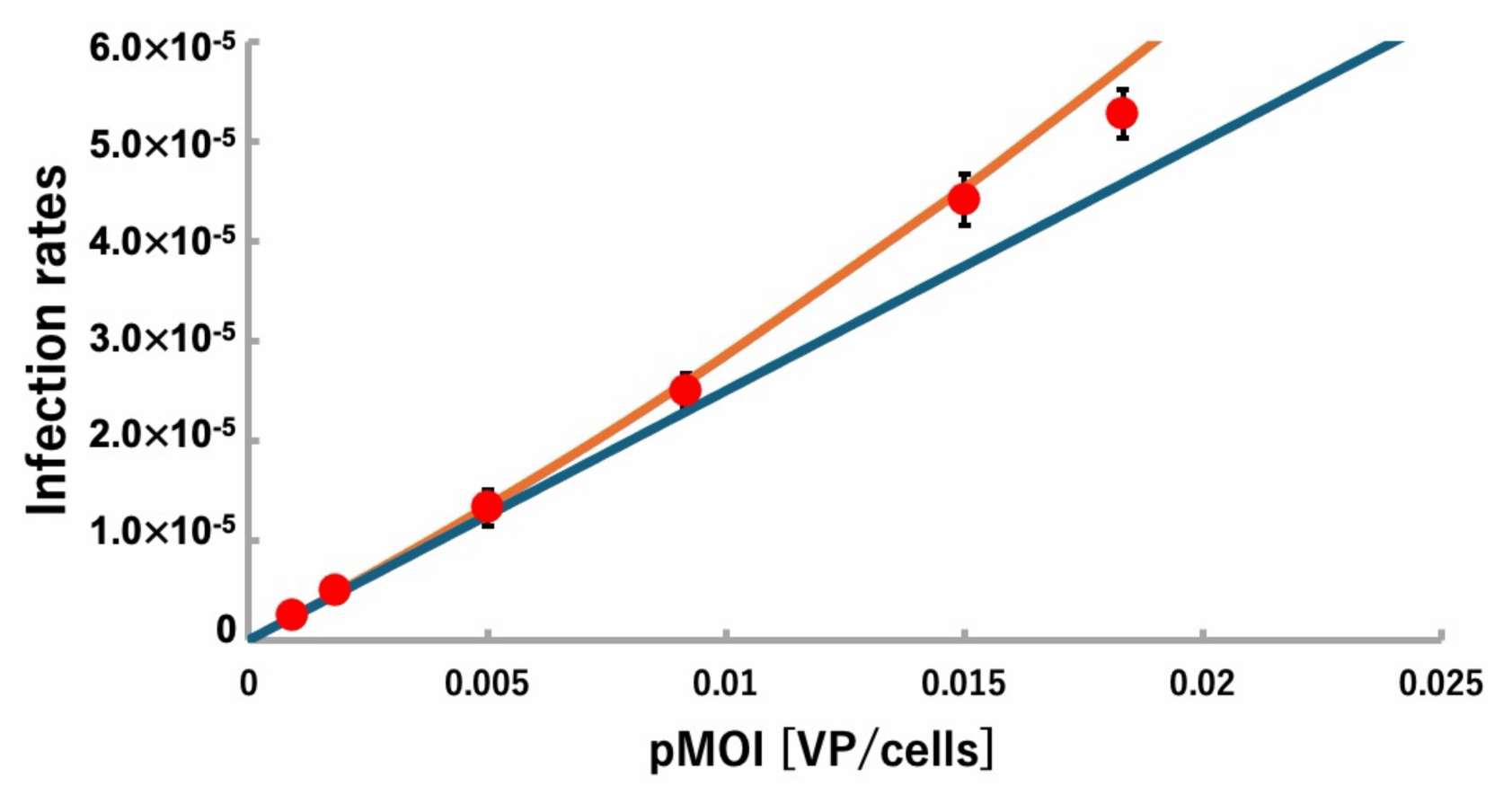
A measurement of infection rates under single-cell conditions and mathematical model of cellular infection. Results of infection rate on low pMOI using plaque assay, in comparison with mathematical models. The x-axis indicates the mean number of virus particles per cell, while the y-axis denotes the cellular infection rate. The red dots in the plot represent experimental data, the orange line represents the model with complementation, and the blue line represents the model without complementation.

### Investigation using different influenza viral strains

To extend our findings with PR8, we next used A/Aichi/2/68 (H3N2) (Aichi) and A/Texas/50/2012 (H3N2) (Texas). Both H1N1 and H3N2 viruses are in circulation among humans on a seasonal basis. Initially, we determined the CTPR for these strains. The obtained CTPR values were then utilized to calculate theoretical infection rates based on a mathematical model of cellular infection. To validate these calculations, actual infection rates were measured under various pMOI conditions at the single-cell level (Fig. 4A, with blue and green lines representing the Aichi and Texas strains, respectively). For the purpose of comparison, the results for the PR8 strain (orange line and red dots) were also included. In both cases, the experimental results reflected the predicted trends of the model. Furthermore, there was a tendency for the number of virus particles required for infection to increase as the CTPR value increased. This suggests that the mathematical model reflects the partial functionality of genome segments within virions, where the number of segments retaining functional integrity varies from particle to particle.

**Fig. 4.**
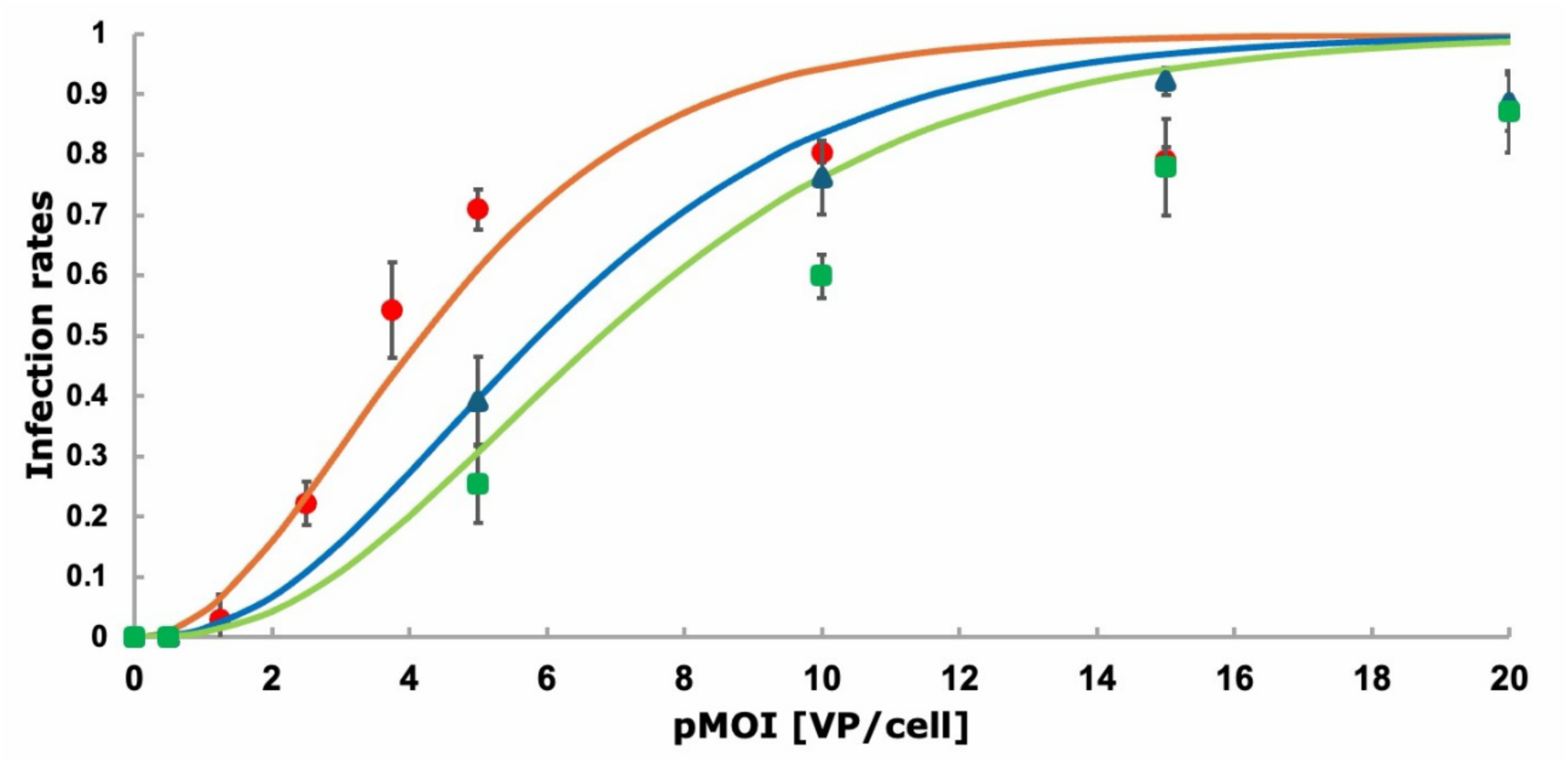
Infection Rate Measurement and Mathematical Model Across Three Influenza Virus Strains. Comparison between infection rate measurements and model predictions for three influenza virus strains: PR8 [Experimental data: red dots, Model: orange line], Aichi [Experimental data: blue dots, Model: blue line], Texas [Experimental data: green dots, Model: green line]. Each model is predicated on the assumption of a complementary infection. The x-axis of this figure corresponds to the pMOI, while the y-axis represents the cellular infection rate.

### Robustness of infection through complementary infection

Furthermore, the CTPR values exhibited a range of more than 20-fold, ranging from 4.0×10² [VP/PFU] for PR8 to 9.2×10³ [VP/PFU] for Texas. However, the pMOI required to achieve approximately 50% infection remained relatively consistent across different strains, ranging from 4 to less than 8 with only a 2-fold variation. To investigate this apparent discrepancy, we conducted simulations of infection rates using our mathematical model while varying CTPR values from 1 to 10^6^ (Fig. 5A). Consistent with the experimental results, the pMOI required to achieve 50% infection remained within the range of approximately 0.5 to 13, even when CTPR values differed by a factor of 10^6^. This suggests that infection can occur even when differences in CTPR is extremely large (even when infectious particles are virtually non-existent). To understand why this occurs, we calculated the distribution of functional genome segments in virus particles based on the *PP* values for the PR8, Aichi, and Texas strains (Fig. 4B–D). These calculations assumed that the number of functional segments per particle follows a binomial distribution, determined based on the number of infectious particles containing eight functional segments (see Methods). Despite the large differences in CTPR values, these distributions differed little. This result suggests that the number of functional genome segments per particle changes only slightly despite the substantial variation in CTPR. Consequently, the number of functional genome segments in the virus particles determines the efficiency of complementation, and complementary infection facilitates efficient infection with a minimal number of virus particles, even in populations containing numerous DVG particles with non-functional segments.

**Fig. 5.**
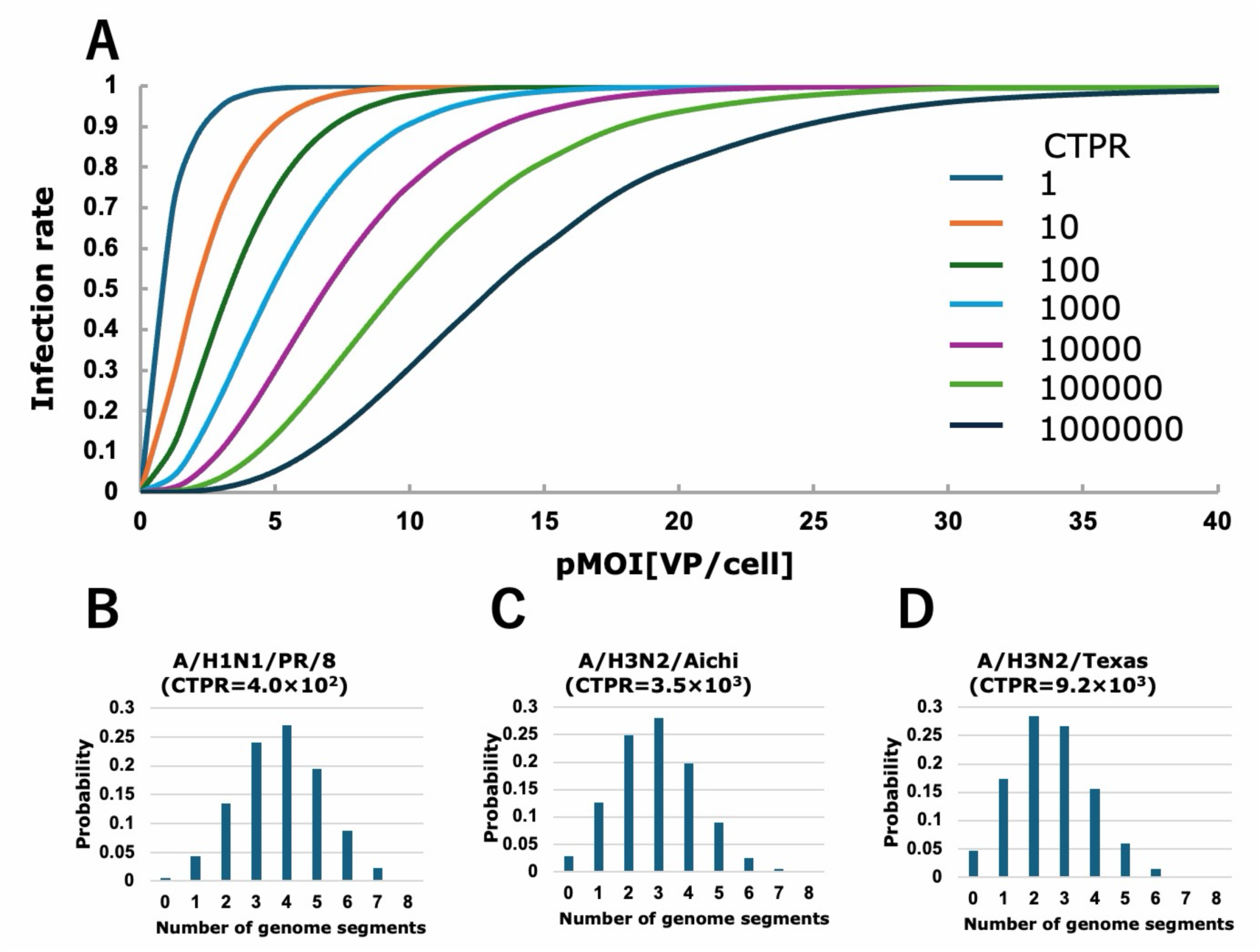
Models of infection rate changes at various CTPR values and predictions of functional genome segment counts within virus particles. (A) Infection rate change models relative to pMOI while varying CTPR from 1 to 10^6^. The calculations were performed according to the multiple infection model used previously in this study. The x-axis represents pMOI, and the y-axis represents the cellular infection rate. (B–D) Distribution of the number of functional genome segments per particle calculated from their respective *PP* values. The proportion of particles containing all eight functional genome segments corresponds to the reciprocal of the CTPR value.

## Discussion

In this study, we performed single-cell infection experiments using quantitatively assessed influenza virus particles under various particle-based multiplicities of infection (pMOI). Our results demonstrated that under high-pMOI conditions, the observed infection rates could not be explained solely by the PFU count. Instead, infection is driven by complementation through simultaneous infection by multiple incomplete virus particles—a classical phenomenon known as multiplicity reactivation ^16,17^. Furthermore, infection experiments using samples with markedly different CTPRs revealed that approximately 4 to 10 physical virus particles are sufficient to achieve 50% cellular infection, regardless of the CTPR value. This trend was corroborated by our mathematical model, which indicated that robust infection occurs under high-pMOI conditions even in a population where only 0.0001% of particles are classically infectious. Biologically, if a virion loses the function of even one of its eight genome segments, it loses its independent infectivity and does not appear as a PFU; however, as long as multiple defective viral genome (DVG) particles retain a subset of functional segments, they can collectively establish infection. While the concept of multiplicity reactivation has long been recognized, the fundamental novelty of this study lies in providing absolute quantitative proof of this phenomenon: we quantitatively demonstrate that multiplicity reactivation is the primary driver of high-pMOI infection, redefining the true infectious unit of influenza A virus as a collective population of particles.

Additionally, the probability that virus particles contain all eight types of intact genome segments was the focus of our research, and a mathematical model was constructed under the assumption of complementation. When comparing the model predictions with experimental results for three influenza virus strains [PR8, Aichi, Texas], the experimental infection rates closely matched the model assuming complementation (Fig. 2D, Fig. 3A, and Fig. 4A). However, the experimental infection rates saturated at approximately 80%, even under conditions of high pMOI. We hypothesize that this suppression at elevated pMOI is caused by defective interfering (DI) particles. It is crucial to strictly distinguish between generally defective particles—which simply lack functional segments—and true DI particles. Within the vast and diverse population of defective particles, DI particles represent a specific subset harboring genome segments with internal deletions but conserved termini, conferring active interfering properties. In our current mathematical model, non-functional segments were treated simply as being absent during the calculation of complementation. Biologically, however, these specifically structured DI segments can be preferentially replicated inside cells. It has been well established that DI particles can suppress the cytopathic effect (CPE) induced by standard virus replication, as evidenced by the studies of Manzoni T. B et al. and Bora M. et al. ^30,31^. Therefore, at extremely high pMOI, the increased probability of co-infecting cells with this specific DI sub-population likely suppresses visible infection markers, capping the observable infection rate.

The mathematical model posits that each of the eight genome segments is randomly packaged into virus particles as either a functional or defective segment. The experimental results obtained in this study align closely with the proposed model, thereby substantiating its validity. However, it is understood that the packaging of the influenza virus genome entails specific interactions among RNA segments designated as packaging signals, resulting in a “1+7” arrangement ^32–34^. In an artificial virus system devoid of the HA segment, it was reported that host-derived 18S and 28S ribosomal RNAs were incorporated as the eighth RNP to preserve the 1+7 structure, thereby underscoring the significance of incorporating eight RNPs into the virus particle ^5^. Consequently, while we acknowledge the established consensus that influenza virions selectively package one copy of each of the eight distinct genome segments, our model and experimental results demonstrate that the functional integrity of these packaged segments is not absolute. Rather, whether each individual segment remains functional follows a stochastic distribution. This indicates that, although the packaging process strictly ensures the inclusion of all eight segment types, the actual functionality of each segment within the cell is governed probabilistically.

On the other hand, the results of this study revealed that the quantification of virus particle numbers has a significant impact on cell infection. In this study, we quantified virus particle numbers using a digital influenza assay ^27^ that we previously developed. This method involves a device containing approximately 800,000 microchambers; it stochastically traps a single influenza viral particle within each chamber and detects fluorescence based on the neuraminidase activity of the influenza virus. In other words, a fluorescent microchamber indicates the presence of a single influenza virus particle, and the virus particle concentration can be determined by dividing this total count of fluorescent microchambers by the total volume of the observed microchambers^27,39^. Since this detection method uses neuraminidase activity as an indicator, there is a possibility of detecting neuraminidase from sources other than whole virus particles or fragments of virus particles. In such cases, the measured value would exceed the actual number of virus particles, resulting in an overestimated CTPR. Consequently, the pMOI value required for infection would also be overestimated. However, we believe that the impact of neuraminidase activity originating from viral fragments or other non-particle sources is limited in the present results. For instance, in the experiment shown in Fig. 2D, where the measured CTPR was 400, if half of the detected entities were non-infectious viral fragments, the true CTPR would actually be 200. If such completely non-infectious viral fragments were present as contaminants, the pMOI would be overestimated, causing the entire experimental plot in the pMOI vs. infection rate plot to shift toward higher pMOI values (Supplement Fig. 1). Conversely, the theoretical model represents ideal infection rates under the assumption that no non-infectious fragments are present. Therefore, the close alignment between the theoretical model and the experimental results strongly implies that contamination by non-infectious entities was minimal. Although false-positive signals originating from non-particle neuraminidase activity may still have been detected to some extent in our experiments, we conclude that their proportion was not substantial.

Conventionally, the majority of studies have examined infection dynamics exclusively in terms of PFU concentrations. The present study underscores the significance of taking virus particle numbers into account, particularly in scenarios involving multiple infections. In practical terms, when considering transmission through respiratory droplets, previous reports indicate that 400 to 5,000 droplets ranging from 0.6 µm to 2,000 µm are generated per cough ^35^, and approximately 1 to 100 PFUs can be detected per cough ^36,37^. This finding indicates that the number of infectious particles, as represented by PFU, present in a single droplet ranges from approximately 0.0025 to 0.02. However, the results of this study suggest that the total number of viral particles per droplet is likely to be 10 to 1,000 times higher, with a maximum of approximately 2.5 to 20 viral particles per droplet. It is noteworthy that, based on the findings of the present study, the minimum number of virus particles required for infection ranges from 4 to 10. This suggests that many of the droplets expelled during coughing and sneezing are infectious. In summary, while the concept of multiplicity reactivation by defective particles has been recognized for over half a century, its quantitative reality has long remained elusive. In this study, by determining the absolute number of physical virus particles, we have finally unveiled the true nature of multiplicity reactivation. Our quantitative approach demonstrates that genome complementation is not merely an auxiliary mechanism, but rather the fundamental driver of infection dynamics. Ultimately, this quantitative unraveling of multiplicity reactivation redefines the true infectious unit of the influenza virus and establishes a new definitive foundation for assessing viral transmission and risks.

## Methods

### Cells and viruses

Madin-Darby Canine Kidney (MDCK) cells were obtained from the American Type Culture Collection. The MDCK cells were cultivated at 37°C with 5% CO2 in Dulbecco’s modified Eagle’s medium (DMEM), which was supplemented with 10% fetal bovine serum (FBS) and 1% penicillin–streptomycin. The preparation of the influenza viruses [A/Puerto Rico/8/1934 (H1N1), A/Aichi/2/1968 (H3N2), A/Texas/50/2012 (H3N2)] was conducted by first confluent MDCK cells on a dish plate, which were then directly infected with a stock sample of each virus at a low multiplicity of infection (MOI) condition in the presence of trypsin. 72 hours after infection, the culture medium’s upper layer was subjected to a centrifugal process at an angular velocity of 1700× *g* for a duration of 10 minutes at a temperature of 4℃. The resultant upper layer was then stored in a cryogenic freezer set at a temperature of −80°C. Prior to utilization, the stored sample was subjected to a thawing process, followed by a dilution procedure to ensure its suitability for use.

### Viral quantification

The digital influenza assay was performed in accordance with the established protocol, as previously described for the measurements of the number of viral particles ^27^. The plaque assay was performed to quantify the number of plaque-forming units. In the case of influenza viruses, the number of plaques was enumerated two days after infection.

### Single cell assay

Sievewell^TM^ chips (Tokyo Ohka Kogyo Co. Ltd.) with 370,000 microwells (20 µm x 25 µm, diameter x depth) were utilized for the observation of individual cells. The chips were meticulously primed with 100% ethanol and phosphate-buffered saline (PBS), followed by thorough washing with the viral culture medium on two separate occasions. The MDCK cells were infected with each virus and subsequently incubated at 35°C to facilitate viral entry. One hour after the infection, the infected cells were loaded into the chip and subsequently incubated at 35°C for 48 hours following the viral infection. The cellular images were captured by the CM30^TM^(EVIDENT), the incubation monitoring system, with two acquisitions: initially upon cell loading into the device and subsequently 48 hours after infection. As a result of viral budding, infected cells undergo the cytopathic effect (CPE) due to viral budding ^38^. This phenomenon leads to the disintegration of the cells’ structural integrity, resulting in their loss of shape and subsequent disappearance from the visual field within the well. Therefore, we painted live and dead cells on the captured images and provided them to the machine as teacher data for learning, which can be adapted to other images after the neural network formation (Supplemental Fig.1). The number of live cells was enumerated, and subsequently, cellular death rates were calculated by the following formula (where *x_i_* denotes the cellular death rate at *MOI* = *i*).

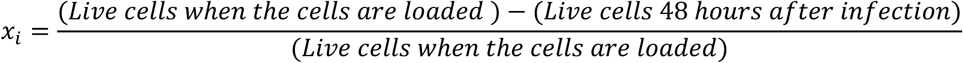

Due to the inherent cellular heterogeneity, a proportion of the cells were found to be dead even in the absence of viruses (*x*_0_ = 0.048). Then, cellular infection rates were calculated using the following formula.

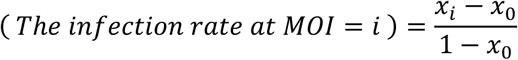

### Mathematical model of viral infection

Mathematical models of viral infection with and without complementation were constructed.

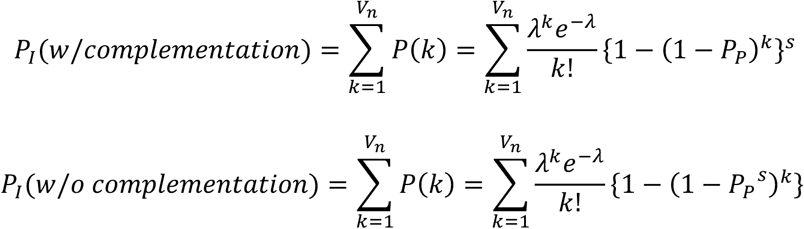

The model was developed under the assumption that the conditions corresponded to a single-cell assay. Consequently, the number of cells (*C_n_*) was designated as 5.0 × 10^4^. The term “*λ*” denotes the expected value of the Poisson distribution and is equivalent to pMOI (= *V_n_*/*C_n_*). Therefore, the number of virus particles (*V_n_*) is expressed as *λC_n_*.

### Distribution of viral functional genome segments

The *P_P_* values were determined through the utilization of the CTPR, subsequently, the distribution of viral functional genome segments was ascertained by employing the following formula:

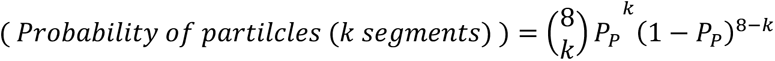

## Supporting information

Simulation of infection rates in the presence of non-particle neuraminidase during virus particle quantification.

## Acknowledgements

This research was supported by the Japan Science and Technology Agency for Core Research for Evolutional Science and Technology, JST CREST, Japan (JPMJCR22N2), and The 2024 - 2026 UTOPIA AI Research Discovery Program, AMED, Japan (JP223fa627001).

## Author Contributions

Yuu K. and K. V. T. designed the study and wrote the paper. Yuu K. and K. T. performed the experiments and analyzed the data. Y.F. and Y.O. performed initial infection experiments using immunostaining and discussed research ideas. S.Y. supported virus quantification by plaque assay and provided critical discussion. Yoshihiro K. and H. N. provided critical discussion and comments.

