## Supplementary material for "The infectious unit of influenza A virus is a population of particles, not a single virion, as shown by absolute particle quantification": Simulation of infection rates in the presence of non-particle neuraminidase during virus particle quantification.

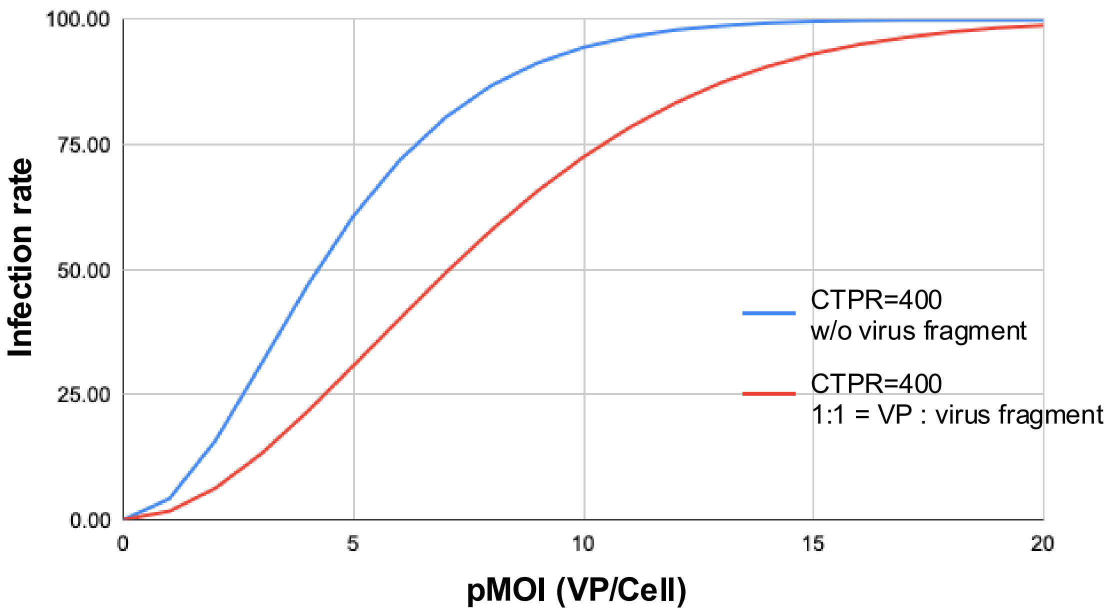


Supplement Fig. 1 Simulation of infection rates in the presence of non-particle neuraminidase during virus particle quantification.

The horizontal axis represents the pMOI determined from virus particle quantification, while the vertical axis represents the infection rate. The blue line indicates the scenario in which no non-particle neuraminidase was included in the virus particle quantification, with all detected signals originating from virus particles. The red line indicates the scenario in which virus particles and non-particle neuraminidase were detected at a 1:1 ratio during the virus particle quantification. Simulations for both scenarios were performed using a CTPR of 400.
